# LocusFocus: A web-based colocalization tool for the annotation and functional follow-up of GWAS

**DOI:** 10.1101/2020.01.02.891291

**Authors:** Naim Panjwani, Fan Wang, Cheng Wang, Gengming He, Scott Mastromatteo, Allen Bao, Jiafen Gong, Johanna M Rommens, Lei Sun, Lisa J Strug

## Abstract

Genome-wide association studies (GWAS) have primarily identified trait-associated loci in the non-coding genome. Colocalization analyses of SNP-level associations from GWAS with expression quantitative trait loci (eQTL) evidence enable the generation of hypotheses about responsible mechanism, genes and tissues of origin to guide functional characterization. Here, we present a web-based colocalization browsing and testing tool named LocusFocus (https://locusfocus.research.sickkids.ca). LocusFocus formally tests colocalization using our established *Simple Sum* method to identify the most relevant genes and tissues for a particular GWAS locus in the presence of high linkage disequilibrium and/or allelic heterogeneity. Full documentation and source code for LocusFocus are publicly available.

## Background

The majority of disease-associated variants identified by genome-wide association studies (GWAS) lie in non-protein-coding regions of the genome [1, 2]. Thus, understanding the mechanism of these non-coding variants and determining which are the causal variant(s) are challenging; linkage disequilibrium (LD), allelic heterogeneity at a given locus, and the polygenic nature of complex traits complicate this effort [3].

Non-coding GWAS variants may tag *cis*-regulatory elements that impact gene expression [4], offering hypotheses on underlying mechanisms that ultimately influence disease phenotype. Integrating GWAS summary statistics with other functional datasets such as expression quantitative trait locus (eQTL) studies, with the aim of understanding what underlying characteristics are driving a particular GWAS locus, is an integral next step post-GWAS necessary to guide functional studies.

Several summary statistic-based colocalization methods are in use, such as coloc [5], eCAVIAR [6, 7], RTC [8], Enloc [9], and moloc [10]. Common challenges for these tools include 1) the impact of LD, 2) allelic heterogeneity [3], and 3) the absence of causal variants in the dataset (untyped or not called), as reviewed in [3].

The *Simple Sum (SS*) [11] is a frequentist colocalization method that, unlike the aforementioned methods, leverages LD information without requirement for causal variant assumptions or prediction. *SS* is more powerful for colocalization than existing methods, especially in regions of high LD and allelic heterogeneity [12]. When integrating an eQTL dataset, the *SS* method not only determines whether a GWAS signal is driven by expression variation but also prioritizes the most probable responsible gene(s) and tissue(s) at the locus. In our previous work on a GWAS of meconium ileus (MI) [11], an intestinal obstruction phenotype in individuals with cystic fibrosis (CF), we showed the information gained through application of the *Simple Sum* colocalization method to genome-wide significant loci with GTEx eQTLs. The *SS* guided the identification of the likely responsible gene(s) for each genome-wide significant locus and pointed to the pancreas as a common contributor in the pathophysiology of MI, a CF phenotype that manifests in the intestine. For example, the genome-wide significant signal detected around the *ATP12A* gene clearly showed colocalization with GTEx eQTLs of *ATP12A* in the pancreas (see figure 2C in [11]), and only the *SS* colocalization method (compared to COLOC and eCAVIAR) highlighted the colocalization, while no support for other digestive system tissues was evident (see Table 3 in [11]). Here we make visualization and testing of colocalization via the *SS* method accessible via a web application named LocusFocus (https://locusfocus.research.sickkids.ca).

LocusFocus allows the user to upload GWAS summary statistics, along with any other secondary SNP-level summary statistic dataset (e.g. eQTL, mQTL and or other phenotypic associations from GWAS) to test colocalization of the two datasets at a particular locus. The user may upload their own sample-specific LD matrix or select the appropriate 1000 Genomes [13] population. LocusFocus guides the functional follow-up in the most probable tissues and genes when used with eQTL datasets. In the examples shown, the primary dataset is a GWAS locus for MI, and the secondary datasets are eQTL p-values from GTEx [14] or those from our own study of primary nasal epithelia from individuals with CF. We have made eQTL summary statistics from GTEx (v7) available for selection within our web server to easily test colocalization with GTEx tissues and genes using the *SS* method.

## Application Design and Implementation

LocusFocus is a web application which uses Python’s Flask as the underlying app engine for *SS* colocalization analysis and subsequent visualization. GTEx v7 [14] eQTL summary statistics for all 48 tissues are indexed and stored in a MongoDB database to enable efficient querying. The plots employ Plotly.js (https://plot.ly/javascript/) to enable interactivity with the data. Figure 1 shows the LocusFocus web interface and required input fields. After submission of the user’s GWAS summary data, colocalization analysis is first performed and colocalization and heatmap plots are then generated for the visualization of GWAS, eQTL, and optional secondary datasets uploaded by the user (a detailed example is shown in Figure 2 for illustrative purposes). Calculation of the LD matrix based on the 1000 Genomes (phase 1, version 3) [13] is performed for the user-specified region and SNPs using PLINK v1.90b6.9 [15]. Alternatively, the user may input their own population-specific PLINK-generated LD matrix for more accurate computation of the *SS* statistic [11]. Users’ integrated data are stored in sessions with unique identifiers for easy sharing of the session data and plots (user sessions are currently stored at least 7 days). The gene track shown below the plots (Figure 2a) is from GENCODE v19 [16], customized to collapse transcript isoforms into single gene models; the gene coordinates were downloaded from GTEx’s web portal. Our current version supports hg19 coordinates; a future version will provide the option to work with hg38 coordinates.

**Figure 1.**
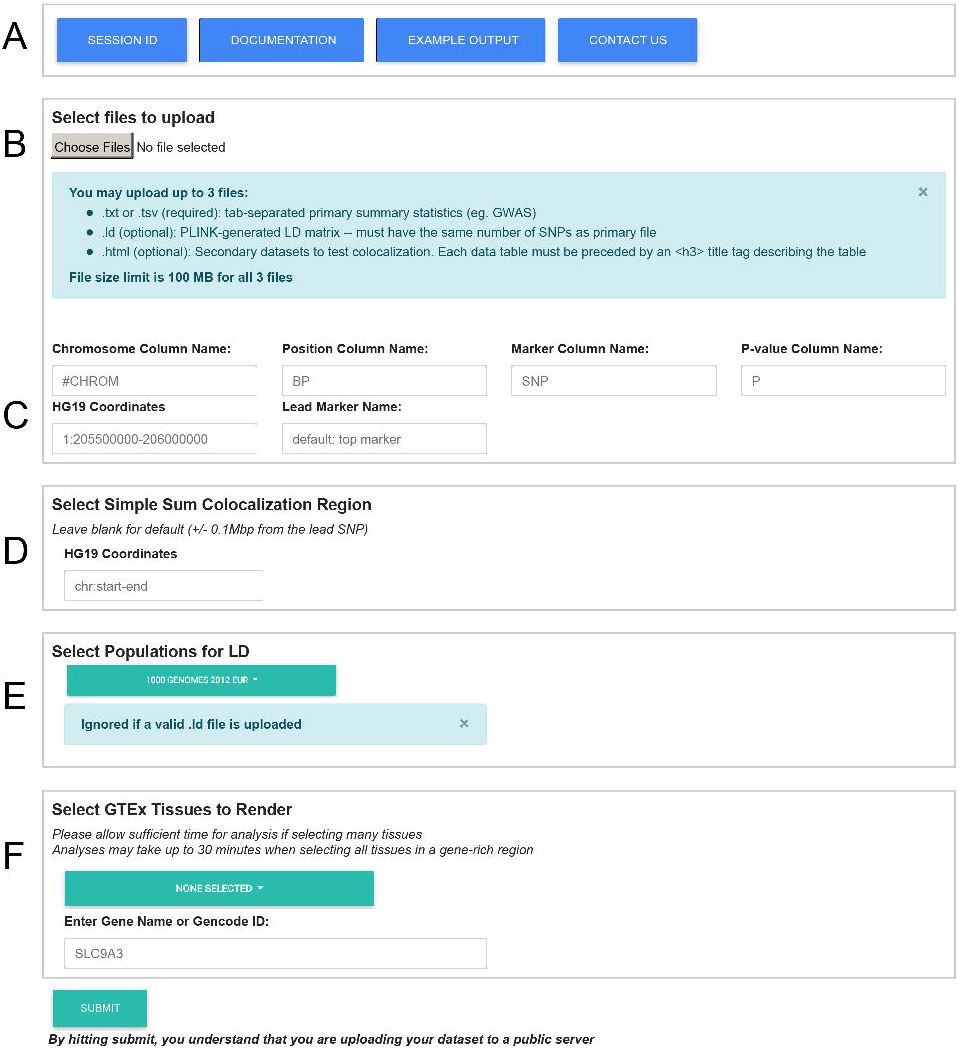
LocusFocus Web Application Input Form. A web-based input form is presented to the user to upload datasets for colocalization analysis at https://locusfocus.research.sickkids.ca. **A.** The *Session ID* button allows the user to retrieve previous colocalization analyses. These sessions are currently stored for at least 7 days. Easy navigation to documentation and example output is provided. **B.** An upload button is provided for up to 3 files not exceeding 100 MB in total (at least the first file is required). File extensions dictate the type of file uploaded: 1) .txt and .tsv files are assumed to be summary statistics for the primary dataset to test colocalization with and is required; this is usually a GWAS dataset. Optionally, one may upload 2) the LD matrix output from PLINK (--r2 square; .ld file extension) and or 3) a multi-sample dataset formatted in HTML format with the secondary summary statistics at the same locus as the primary dataset to test colocalization with. **C.** Column names for the primary dataset may be changed here. Four columns, in any order, are required (chromosome, position, marker column name with rsid or chrom_pos_ref_alt_b37, and a p-value column), while other columns are ignored. The user may enter the column names for these in the input form. The hg19 coordinates to view plot results are also required (limited to 2 Mbp regions). The lead SNP with the lowest p-value is chosen as default but the user may input alternate lead SNPs. **D.** The *Simple Sum* or *SS* region tests colocalization across a default region of 0.1 Mbp on either side of the lead SNP, but the user may input a customized region up to 2 Mbp (the evaluated area will appear in gray shading in the first plot output). **E.** Can be ignored if a user .Id file was provided in B, otherwise, the 1000 Genomes population [13] that most closely resembles the input dataset may be selected. **F.** Secondary datasets from any, any subgroup or all 48 tissues from GTEx (v7) [14] are available for selection within the webserver. An initial gene input is required for the initial colocalization plot across all tissues selected, but the eQTLs for all genes that fall within the region provided in **C** are tested for colocalization and each gene is made available for plotting in the subsequent output page via a dropdown. Colocalization is tested for each of the tissues selected and for all genes in the input region **C**.

**Figure 2.**
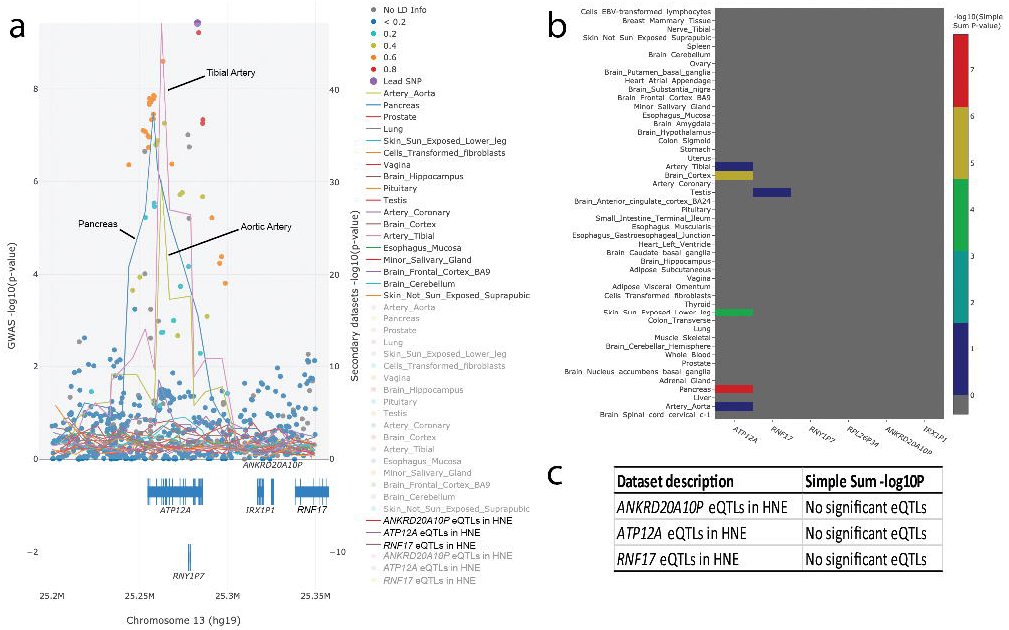
Sample interactive plot output from the LocusFocus web application. GWAS summary statistics of MI and lung disease severity in patients with cystic fibrosis for chr13q12.12 were uploaded, and all tissues from GTEx [14] were selected for colocalization analysis. The plots are traced using plotly in JavaScript (https://plot.ly/javascript/) after merging of the input data. **a)** Filled circles represent GWAS data (with corresponding y-axis on the left) for MI. The LD information presented is similar to LocusZoom [26] (lead SNP in purple, high LD SNPs with r^2^ ≥ 0.8 in red with the lead SNP, orange for 0.8 < r^2^ ≤ 0.6, green for 0.6 < r^2^ ≤ 0.4, light blue for 0.4 < r^2^ ≤ 0.2 and dark blue for r^2^ < 0.2; markers with no LD information are shown in gray). LD information was computed from the European 1000 Genomes subset (phase 1, version 3) [13]. The web server computes the LD matrix with the 1000 Genomes on demand using PLINK v1.90b6.9 [15]. Lines shown on the plot represent a summary of GTEx (v7) and primary human nasal epithelial cells’ (HNEs) eQTL p-values for *ATP12A,* a gene proposed as a modifier for CF [17–19] (with corresponding y-axis on the right), with each line representing a tissue (eQTLs for other genes within the region can be selected and the plot re-drawn within the same session). Line traces for some tissues do not appear due to no eQTL data for *ATP12A* for that tissue (likely due to little or no expression). The lines trace the lowest p-value per window, and the windows are defined as (region size/1,000,000) × 150, where region size is the size of the region input in base pairs (up to 2 Mbp regions are allowed). A different window size can be specified and lines redrawn on the web tool. We find these parameters best illustrate the overall pattern of eQTL association for a particular window size up to 2 Mbp. Gene track information is from GENCODE v19 (hg19 coordinates), with transcripts collapsed into single genes (as described by GTEx). The gray shaded region shows the region used for the *SS* calculation, 0.1 Mbp on each side of the selected lead SNP (by default unless set differently by user). We used the full region (chr13:25,200,000-25,350,000) for the *SS* calculations. Users may click the tissue panel list in the legend to show or hide particular groups of information. The eQTL scatterplots for each tissue, from which the line traces are derived from, are hidden by default but may be displayed by clicking on the desired tissue in the legend (tissues listed in faint gray; not all tissues are shown in the colocalization figure above due to space; for a complete table of colocalization results, refer to Supplementary Table 1, or view interactively at https://locusfocus.research.sickkids.ca/session_id/e688fad4-0746-4d71-8730-15cd5e4b12bf or bit.ly/LocusFocus-Example). Other features of the plot include the ability to zoom in, tooltips for each data point, save image options in png format, selection and fading tools, and resetting, rescaling or shifting of axes. The heatmaps shown in lower panels summarize the *SS* colocalization tests for all the genes in the user-defined region and across all the selected tissues. Gray squares with negative p-values for colocalization indicate either no eQTL data (typiclally due to little or no expression), or the gene-tissue pair does not have significant eQTL signal after Bonferroni correction, or insufficient SNPs are provided for an accurate calculation of the *Simple Sum* p-value (the exact reason can be viewed in the web session as an interactive table output, or in Supplementary Table 1).

All code is publicly available via GitHub (https://github.com/naim-panjwani/LocusFocus) under the MIT license, providing a means to install LocusFocus in a local web server.

## Colocalization Analysis with LocusFocus Near *ATP12A*

We illustrate the use of LocusFocus for the chr13q12.12 association around *ATP12A* from the GWAS of MI in CF [11]. As stated earlier, we found strong colocalization of *ATP12A* in the pancreas but no colocalization with other nearby genes and digestive system tissues in [11]. The GWAS summary statistics for MI at the chr13q12.12 (chr13:25,200,000-25,350,000; hg19) locus were uploaded into LocusFocus, and the 1000 Genomes [13] European population was chosen for the LD matrix calculation (Figure 1). The GWAS summary statistics, LD matrix and eQTL data are integrated with outputs of interactive colocalization and heatmap plots (Figure 2), and interactive summary tables (Figure 2c and Supplementary Table 1). Results support strong colocalization for *ATP12A* in the pancreas as reported [11]. Interestingly, *ATP12A* has been proposed as a modifier of lung disease severity via its role in pH regulation, which in turn may influence immune response to infections in the CF lung [17–19]. Our GWAS on lung disease severity in CF, however, revealed no association at this locus [20]. To further investigate *ATP12A* in the lungs in the event that the CF lung GWAS was confounded, we tested colocalization at the CF MI GWAS locus with eQTLs in lung tissue from GTEx [11] and eQTLs from RNAseq of human nasal epithelia (HNE) from individuals with CF [21]. eQTLs for *ATP12A* from the lung and HNE [22] do not colocalize with the MI GWAS locus (Figure 2). Investigating colocalization of the GWAS p-values with HNE eQTLs for other genes at the locus indicates no significant eQTLs (*ANKRD20A10P*, *ATP12A*, *RNF17*), or a lack of expression in HNE (*RNY1P7*, *RPL26P34*, *IRX1P1*) (Figure 2a, c). Taken together, while biological studies suggest a role for *ATP12A* in the lung [17–19] and have advocated for ATP12A as an alternative CF therapeutic target [23], our studies in the CF population do not support that common variation in expression of this gene affects lung function outcomes.

The current analysis is limited by the tissue sampling, which is confined by source and cell types present. The respiratory epithelium is thought to be relevant tissue in CF lung disease [24]. Lung tissues from GTEx may have limited epithelial components represented and are harvested from individuals without CF [14]. The HNE model addresses these aforementioned concerns, yet temporal- and context-specific expression of any gene are major determinants of function and thus could influence the findings. Although clear evidence of eQTLs for *ATP12A* from pancreas tissue collected by GTEx colocalize with the MI GWAS, this tissue is from adults, while MI is a condition that manifests *in utero* in patients with CF. Colocalization applications clearly benefit from the best secondary datasets available.

## Conclusions

LocusFocus evaluates the colocalization of GWAS with a secondary SNP-level dataset such as eQTLs using a web-based application, enabling the prioritization of the best co-localizing gene(s) and tissue(s) for further follow-up with functional studies. Functionality for secondary datasets with broad (such as GTEx) or more specific relevant data for the traits being analysed is available (e.g. mQTL or other phenotypic datasets). LocusFocus demonstrated our MI GWAS locus at chr13q12.12 colocalizes with eQTLs of *ATP12A* in the pancreas, suggesting the associated variants likely impact MI development through variation in gene expression of *ATP12A* in the pancreas. LocusFocus analysis provided no support for other nearby genes or for *ATP12A* in the lungs.

## Methods

### Documentation and Source Code

Full documentation for LocusFocus is available at https://locusfocus.readthedocs.io. Source code is available at https://github.com/naim-panjwani/LocusFocus.

### Calculation of eQTLs from RNAseq of human nasal epithelia

HNEs were harvested from individuals with CF as described in [21].

RNA-seq data from HNEs was cleaned, aligned and normalized using Trim Galore (v0.4.4), STAR (v2.5.4b), RNA-SeQC (v2.0.0), respectively. The trimmed mean of M values (TMM) method was used. Genotype data from the 610Q, 660W and OMNI 2.5 microarrays underwent individual and SNP-level quality control, and were imputed using a hybrid reference panel consisting of the 1000 Genomes and 101 sequenced individuals with CF [25]. The GTEx (v7) protocol was followed exactly for computation of eQTLs, except the regression model was adjusted with CD45 (PTPRC) normalized expression and the RNA integrity number (RIN).

## Declarations

### Ethics approval and consent to participate

The RNA-seq study was approved by the Research Ethics Board of the Hospital for Sick Children. All CF patients provided informed consent to participate.

### Availability of data and materials

The datasets used, and detailed examples, are available on the LocusFocus GitHub repository (https://github.com/naim-panjwani/LocusFocus). The datasets and file names used are as follows:

- MI GWAS around *ATP12A*: MI_GWAS_2019_13_25180-25400kbp.tsv (subset from the main MI GWAS study [11])
- Secondary HNE eQTL data: slc26a9_uk_biobank_spirometry_merged.html
- GTEx eQTL data: available on the GTEx portal, and made available within our tool
- The session generated has been archived and is available at: https://locusfocus.research.sickkids.ca/session_id/e688fad4-0746-4d71-8730-15cd5e4b12bf or simply bit.ly/LocusFocus-Example

Further examples for the usage of LocusFocus are available as well in the documentation (https://locusfocus.readthedocs.io/en/latest/examples.html).

### Competing Interests

The authors declare that they have no competing interests.

### Funding

This project was supported by Canadian Institutes of Health Research (CIHR MOP 258916, MOP 117978), Cystic Fibrosis Canada (CFC; #2626) and the CFIT Program funded by the SickKids Foundation and CF Canada; Natural Sciences and Engineering Research Council of Canada (NSERC RGPIN-2015-03742, 250053-2013); Genome Canada through the Ontario Genomics Institute (2004-OGI-3-05; 2018-OGI-148); and the US CF Foundation (STRUG17PO).

### Authors’ Contributions

NP designed, implemented and created the LocusFocus web application, and was a major contributor in writing the manuscript. FW developed the *Simple Sum* statistical method, implemented the code for the application, and contributed to the manuscript. CW and GH derived the eQTL calculations from RNAseq of HNEs. SM helped with the implementation and testing of the *Simple Sum* code method. AB helped with testing and identification of implementation bugs. JG helped with the acquisition of data. JMR, LS and LJS defined the application and data analysis, provided guidance in the project and edited the manuscript. All authors read and approved the final manuscript.

## Notes

https://locusfocus.research.sickkids.ca/

https://github.com/naim-panjwani/LocusFocus

