## Supplementary Table 1 for "LocusFocus: A web-based colocalization tool for the annotation and functional follow-up of GWAS"

**Supplementary Table 1.** Simple Sum (SS) colocalization tests for the Genome-wide Association Study (GWAS) of Meconium Ileus (MI) in individuals with Cystic Fibrosis (CF) at the ATP12A (chr13q12.12) locus with all GTEx (v7) tissues, and primary human nasal epithelia (HNE) from individuals with CF. Cell values are -log_10_(SS p-values). Values below were extracted from the LocusFocus web application (session ID e688fad4-0746-4d71-8730-15cd5e4b12bf). Strength of colocalization is coloured from green (low -log_10_P) to red (high -log_10_P). Results support a strong colocalization of ATP12A eQTLs in the pancreas with the GWAS of MI. Gene/tissue cells described as “No eQTL data” (output as -1 by LocusFocus) have no eQTLs calculated by GTEx, likely due little or no expression; “No significant eQTLs” (output as -2 by LocusFocus) describes the scenario where eQTL data is available, but the overall eQTL p-values in relation to other eQTLs does not pass a Bonferroni-corrected threshold prior to SS colocalization testing; a third scenario (which does not occur in this table) for a missing SS p-value is “SS test failed” (output as -3 by LocusFocus), which is often due to an insufficient number of SNPs for a confident assessment of the SS colocalization test.

| **Tissue/Gene** | ***ANKRD20A10P*** | ***ATP12A*** | ***IRX1P1*** | ***RNF17*** | ***RNY1P7*** | ***RPL26P34*** |
| --- | --- | --- | --- | --- | --- | --- |
| **Brain_Spinal_cord_cervical_c-1** | No eQTL data | No eQTL data | No eQTL data | No eQTL data | No eQTL data | No eQTL data |
| **Artery_Aorta** | No eQTL data | 0.626 | No eQTL data | No eQTL data | No eQTL data | No eQTL data |
| **Liver** | No eQTL data | No eQTL data | No eQTL data | No eQTL data | No eQTL data | No eQTL data |
| **Pancreas** | No eQTL data | 7.76 | No eQTL data | No eQTL data | No eQTL data | No eQTL data |
| **Adrenal_Gland** | No eQTL data | No eQTL data | No eQTL data | No eQTL data | No eQTL data | No eQTL data |
| **Brain_Nucleus_accumbens_basal_ganglia** | No significant eQTLs | No eQTL data | No eQTL data | No eQTL data | No eQTL data | No eQTL data |
| **Prostate** | No eQTL data | No significant eQTLs | No eQTL data | No eQTL data | No eQTL data | No eQTL data |
| **Whole_Blood** | No eQTL data | No eQTL data | No eQTL data | No eQTL data | No eQTL data | No eQTL data |
| **Brain_Cerebellar_Hemisphere** | No eQTL data | No eQTL data | No eQTL data | No significant eQTLs | No eQTL data | No eQTL data |
| **Muscle_Skeletal** | No eQTL data | No eQTL data | No eQTL data | No eQTL data | No eQTL data | No eQTL data |
| **Lung** | No eQTL data | No significant eQTLs | No eQTL data | No eQTL data | No eQTL data | No eQTL data |
| **Colon_Transverse** | No eQTL data | No eQTL data | No eQTL data | No eQTL data | No eQTL data | No eQTL data |
| **Skin_Sun_Exposed_Lower_leg** | No eQTL data | 4.07 | No eQTL data | No eQTL data | No eQTL data | No eQTL data |
| **Thyroid** | No eQTL data | No eQTL data | No eQTL data | No significant eQTLs | No eQTL data | No eQTL data |
| **Cells_Transformed_fibroblasts** | No eQTL data | No significant eQTLs | No eQTL data | No eQTL data | No eQTL data | No eQTL data |
| **Adipose_Visceral_Omentum** | No eQTL data | No eQTL data | No eQTL data | No eQTL data | No eQTL data | No eQTL data |
| **Vagina** | No eQTL data | No significant eQTLs | No eQTL data | No eQTL data | No eQTL data | No eQTL data |
| **Adipose_Subcutaneous** | No eQTL data | No eQTL data | No eQTL data | No eQTL data | No eQTL data | No eQTL data |
| **Brain_Hippocampus** | No significant eQTLs | No significant eQTLs | No eQTL data | No eQTL data | No eQTL data | No eQTL data |
| **Brain_Caudate_basal_ganglia** | 0 | No eQTL data | No eQTL data | No eQTL data | No eQTL data | No eQTL data |
| **Heart_Left_Ventricle** | No eQTL data | No eQTL data | No eQTL data | No eQTL data | No eQTL data | No eQTL data |
| **Esophagus_Gastroesophageal_Junction** | No eQTL data | No eQTL data | No eQTL data | No eQTL data | No eQTL data | No eQTL data |
| **Esophagus_Muscularis** | No eQTL data | No eQTL data | No eQTL data | No significant eQTLs | No eQTL data | No eQTL data |
| **Small_Intestine_Terminal_Ileum** | No significant eQTLs | No eQTL data | No eQTL data | No eQTL data | No eQTL data | No eQTL data |
| **Pituitary** | No eQTL data | No significant eQTLs | No eQTL data | No eQTL data | No eQTL data | No eQTL data |
| **Brain_Anterior_cingulate_cortex_BA24** | No significant eQTLs | No eQTL data | No eQTL data | No eQTL data | No eQTL data | No eQTL data |
| **Testis** | No significant eQTLs | No significant eQTLs | No significant eQTLs | 0.0000026 | No eQTL data | No eQTL data |
| **Artery_Coronary** | No eQTL data | 0 | No eQTL data | No eQTL data | No eQTL data | No eQTL data |
| **Brain_Cortex** | No significant eQTLs | 5.4 | No eQTL data | No eQTL data | No eQTL data | No eQTL data |
| **Artery_Tibial** | No eQTL data | 0.833 | No eQTL data | No eQTL data | No eQTL data | No eQTL data |
| **Uterus** | No eQTL data | No eQTL data | No eQTL data | No significant eQTLs | No eQTL data | No eQTL data |
| **Stomach** | No eQTL data | No eQTL data | No eQTL data | No eQTL data | No eQTL data | No eQTL data |
| **Colon_Sigmoid** | No eQTL data | No eQTL data | No eQTL data | No significant eQTLs | No eQTL data | No eQTL data |
| **Brain_Hypothalamus** | No significant eQTLs | No eQTL data | No eQTL data | No eQTL data | No eQTL data | No eQTL data |
| **Brain_Amygdala** | No significant eQTLs | No eQTL data | No eQTL data | No eQTL data | No eQTL data | No eQTL data |
| **Esophagus_Mucosa** | No significant eQTLs | No significant eQTLs | No eQTL data | No eQTL data | No eQTL data | No eQTL data |
| **Minor_Salivary_Gland** | No eQTL data | No significant eQTLs | No eQTL data | No eQTL data | No eQTL data | No eQTL data |
| **Brain_Frontal_Cortex_BA9** | No significant eQTLs | No significant eQTLs | No eQTL data | No eQTL data | No eQTL data | No eQTL data |
| **Brain_Substantia_nigra** | No significant eQTLs | No eQTL data | No eQTL data | No eQTL data | No eQTL data | No eQTL data |
| **Heart_Atrial_Appendage** | No eQTL data | No eQTL data | No eQTL data | No eQTL data | No eQTL data | No eQTL data |
| **Brain_Putamen_basal_ganglia** | No significant eQTLs | No eQTL data | No eQTL data | No eQTL data | No eQTL data | No eQTL data |
| **Ovary** | No eQTL data | No eQTL data | No eQTL data | No significant eQTLs | No eQTL data | No eQTL data |
| **Brain_Cerebellum** | No eQTL data | No significant eQTLs | No eQTL data | No significant eQTLs | No eQTL data | No eQTL data |
| **Spleen** | No eQTL data | No eQTL data | No eQTL data | No eQTL data | No eQTL data | No eQTL data |
| **Skin_Not_Sun_Exposed_Suprapubic** | No eQTL data | No significant eQTLs | No eQTL data | No eQTL data | No eQTL data | No eQTL data |
| **Nerve_Tibial** | No eQTL data | No eQTL data | No eQTL data | No significant eQTLs | No eQTL data | No eQTL data |
| **Breast_Mammary_Tissue** | No eQTL data | No eQTL data | No eQTL data | No eQTL data | No eQTL data | No eQTL data |
| **Cells_EBV-transformed_lymphocytes** | No eQTL data | No eQTL data | No eQTL data | No eQTL data | No eQTL data | No eQTL data |
| **Human nasal epithelia (HNE) from individuals with CF** | No significant eQTLs | No significant eQTLs | No eQTL data | No significant eQTLs | No eQTL data | No eQTL data |
